# Transcriptional plasticity of virulence genes provides malaria parasites with greater adaptive capacity for avoiding host immunity

**DOI:** 10.1101/2024.03.08.584127

**Authors:** Francesca Florini, Joseph E. Visone, Evi Hadjimichael, Shivali Malpotra, Christopher Nötzel, Björn F.C. Kafsack, Kirk W. Deitsch

**Affiliations:** Department of Microbiology and Immunology, Weill Cornell Medical College, New York, New York, USA

**Keywords:** single cell RNAseq, antigenic variation, gene expression, *var* genes, immune evasion

## Abstract

Chronic, asymptomatic malaria infections contribute substantially to disease transmission and likely represent the most significant impediment preventing malaria elimination and eradication. *Plasmodium falciparum* parasites evade antibody recognition through transcriptional switching between members of the *var* gene family, which encodes the major virulence factor and surface antigen on infected red blood cells. This process can extend infections for up to a year; however, infections have been documented to last for over a decade, constituting an unseen reservoir of parasites that undermine eradication and control efforts. How parasites remain immunologically “invisible” for such lengthy periods is entirely unknown. Here we show that in addition to the accepted paradigm of mono-allelic *var* gene expression, individual parasites can simultaneously express multiple *var* genes or enter a state in which little or no *var* gene expression is detectable. This unappreciated flexibility provides parasites with greater adaptive capacity than previously understood and challenges the dogma of mutually exclusive *var* gene expression. It also provides an explanation for the antigenically “invisible” parasites observed in chronic asymptomatic infections.

## Introduction

Despite intensive efforts directed towards malaria eradication, it remains one of the most reported infectious diseases in many tropical countries (1). It is caused by parasites of the genus *Plasmodium*, with *P. falciparum* responsible for the most infections and deaths. These parasites cause disease through asexual replication inside red blood cells (RBCs), generating 20-30 new infectious merozoites every 48 hours. Over the course of their replicative cycle, parasites extensively modify the infected RBC, resulting in more rigid and spherical cells with altered membrane permeability (2). These changes in shape and deformability make the infected cells vulnerable to filtration and elimination by the spleen. To avoid splenic clearance, *P. falciparum* produces and exports several adhesins to the surface of the infected RBCs. *P. falciparum* Erythrocyte Membrane Protein 1 (PfEMP1), the best characterized protein of these adhesins, allows infected RBCs to attach to the vascular endothelium by binding tissue specific receptors, thereby enabling infected cells to sequester from the peripheral circulation and avoid passage through the spleen (3–5). This cytoadhesion and sequestration of infected RBCs leads to formation of RBC aggregates within capillary beds and can result in the interruption of blood flow and localized inflammation that underlies the severe syndromes of cerebral and placental malaria (6–8).

While the vast majority of parasite produced proteins are hidden within the infected RBC, the exposure of PfEMP1 on the infected cell surface makes it the primary target of the humoral immune response (9). Antibodies against PfEMP1 can greatly reduce parasitemia, however, parasites can escape complete elimination by exchanging the PfEMP1 isoform expressed, thereby enabling small populations of parasites to expand and perpetuate the infection in a process called antigenic variation (10). This is possible because different PfEMP1 isoforms are each encoded by individual members of the *var* multicopy gene family comprised of 45-90 paralogs per haploid genome (11, 12). *var* gene expression is thought to be mutually exclusive, meaning that only one gene is transcriptionally active at a time, and by switching expression from one gene to another, parasites can change the PfEMP1 displayed on the infected cell surface, leading to the waves of parasitemia that are characteristic of a *P. falciparum* infection. Transcriptional control is epigenetically regulated through the deposition of specific activating and silencing histone marks, leading to condensed heterochromatin formation at all *var* loci except the single transcriptionally active copy (13–15). This process directly links parasite virulence to the persistent nature of malaria and is critical for the parasite’s ability to survive, cause disease and be efficiently transmitted.

Antigenic variation through *var* gene switching is thought to enable an infection to persist for a year or longer as the parasite population cycles through its repertoire of *var* genes (16, 17). However, a puzzling aspect of *P. falciparum* infections is the occasional identification of chronic, asymptomatic infections that can last for a decade or more (18). In such instances, it is presumed that the *var* gene repertoire would have been exhausted, leaving unexplained how the parasites have avoided complete elimination. One detailed study of an asymptomatic infection that was only detected upon splenectomy identified parasites that were no longer expressing *var* genes and had lost any measurable cytoadhesive properties (19). Such parasites could potentially be immunologically “invisible” due to a lack of surface antigen expression, thus explaining their lengthy persistence as well as their resurgence upon removal of the spleen. These parasites regained *var* gene expression when subsequently grown in culture, indicating that this “*var*-null” state was a selected phenotype rather than resulting from mutation. Moreover, the loss of *var* gene expression suggests the possibility of greater flexibility in how this gene family is regulated. However, how common it is for parasites to completely silence *var* gene expression or how this could fit into a molecular mechanism that typically ensures single *var* gene expression is unknown.

Mutually exclusive expression is a well-conserved mechanism displayed by organisms throughout the eukaryotic evolutionary tree, including a number of eukaryotic pathogens for immune evasion, for example *Trypanosoma brucei* (20, 21), *Giardia lamblia* (22) and *Babesia bovis* (23). It is also evident in the expression of the immunoglobulin and olfactory receptor genes in mammals (24, 25). In the mammalian examples, mutually exclusive expression is part of terminal cellular differentiation and thus the choice of which gene is activated is permanent. In contrast, for parasites that employ this process for antigenic variation, the choice of which gene is activated is semi-stable and reversible, enabling parasites to repeatedly activate and silence different genes over the course of an infection, adding another layer of complexity to an already poorly understood phenomenon. Decades of research on mutually exclusive gene expression has historically relied on information obtained from populations of cells. Recently, advances in single-cell-based approaches have begun to reveal that cells within a population can be very heterogeneous, and that these differences can be revealing for understanding the molecular mechanisms behind various biological processes. Only through single-cell methodologies has it been possible to begin to decipher the complexity of immunoglobulin choice during lymphocyte development (26, 27) or the maturation of olfactory neurons that results in expression of a single olfactory receptor (28–30).

To date, several studies have successfully applied single-cell RNA-Seq to *Plasmodium falciparum* and uncovered crucial signatures necessary for processes involved in replication, life-cycle progression, and transmission (31–33). However, no study has yet specifically focused on *var* gene expression, despite the importance of these genes for parasite survival and pathogenesis. We recently approached the question of *var* gene choice through the analysis of an extensive library of closely related, recently isolated clonal parasite populations (34). We observed two distinct *var* expression profiles: either high level expression of a single *var* gene, as expected for a population of parasites exhibiting mutually exclusive expression of a single dominant *var* gene, or alternatively low-level expression of a large portion of the *var* gene family, an unanticipated state that is of unknown significance. In addition, parasites were able to switch between these two states, suggesting that this observation could provide clues to how mutually exclusive expression and *var* gene switching might be regulated. However, in the absence of single cell resolution, it was not possible to understand the molecular mechanisms underlying these distinct *var* expression profiles or the possible significance of this phenomenon.

In the present study, we focused on understanding how individual cells contribute to the cumulative *var* expression profile of a parasite population. We combined different single-cell RNA-Seq approaches to analyze *var* gene expression in recently cloned lines at single-cell resolution. Our results demonstrate that individual parasites can exist in three distinct *var* expression states. In addition to the expected parasites expressing one *var* gene at a high level, we identified parasites expressing more than one *var* gene simultaneously and parasites in which *var* gene expression was barely detectable and very heterogenous. Manipulation of intracellular S-adenosylmethionine levels, the methyl donor required for histone methylation and heterochromatin formation, enabled us to skew parasites toward different *var* expression states, providing clues to the molecular mechanisms underlying single *var* gene choice. The discovery that parasites have much greater flexibility in *var* gene expression, including the ability to silence PfEMP1 expression and become antigenically “invisible”, significantly changes our understanding of how *P. falciparum* undergoes antigenic variation and has important implications for elimination of chronic malaria infections.

## Results

### Clonal parasite populations from different genetic backgrounds display two states of *var* expression

The well-accepted, standard paradigm regarding malaria antigenic variation assumes that at any given time, individual parasites express one and only one member of the *var* gene family. In recent work from our laboratory, we generated an extensive collection of closely related, isogenic 3D7 and NF54 subclones through limiting dilution, thereby enabling us to examine recently cloned parasite populations expressing alternative *var* genes (34). To our surprise, not all the clones displayed the predicted pattern of single *var* gene expression. Analysis by quantitative real-time RT-PCR (qRT-PCR) and RNA-Sequencing of the different populations showed that while some lines were expressing high levels of a single *var* gene, as predicted by a model of mutually exclusive expression, others displayed very low-level expression of a heterogenous mix of *var* genes. While previous studies similarly reported substantial differences in levels of *var* gene expression in recently subcloned lines of various strains (35, 36), we nonetheless wanted to verify that this was not an artifact of the prolonged in vitro culturing of the 3D7 strain. We therefore generated a new collection of isogenic subclones from a population of the Asian isolate IT4 (also called FCR3) (37). Using *var* specific qRT-PCR, we analyzed populations obtained from two consecutive rounds of subcloning and determined the *var* expression profiles. Similar to 3D7, we detected parasites either expressing a single *var* gene at a high level (called the “high-single” expression state) or very low levels of expression of multiple *var* genes (referred to as “low-many” populations) (Supplemental Figure S1), indicating that this phenomenon is not unique to 3D7.

Measurement of cumulative *var* expression from each of the analyzed IT4 clones showed that parasites in the “low-many” state displayed significantly lower total *var* expression, similar to what we previously observed for the 3D7 and NF54 strains (34). These data strongly suggest that *P. falciparum* parasite populations do not always express a dominant *var* gene, as previously assumed. However, this analysis as well as previously published research on *var* expression were performed on populations of parasites encompassing millions of individual infected cells. This poses an obvious limit to our interpretation of the data as it obscures the contribution of individual parasites to the cumulative profile. In the specific case of the clonal populations in the “low-many” state, it is impossible to distinguish if individual parasites are expressing a normal level of *var* gene transcripts but rapidly switching between *var* genes, thus resulting in the lack of a detectable dominant gene, or if all the individual parasites are actually in a “low-many” state.

### Analysis by single-cell RNA-Seq of “high-single” and “low-many” populations

To examine *var* gene expression in individual parasites, we analyzed different wildtype, recently cloned 3D7 populations by Drop-Seq, a droplet-based single-cell RNA-Seq (scRNA-Seq) method previously adapted to *P. falciparum* (38). Since our interest is focused on *var* genes, which are expressed between 10 and 20 hours after merozoites enter the red cell, we tightly synchronized the populations and isolated parasites at 16-19 hours post-invasion (hpi). We sequenced cDNA from individual infected cells from two recently cloned populations in the “high-single” state, expressing one *var* gene at high levels, and two populations in the “low-many” state, expressing low-levels of a heterogenous mix of *var* genes. To confirm the *var* expression status of the populations when individual infected cells were isolated, *var* profiles were assessed by qRT-PCR on total RNA extracted from the population on the same day that the scRNA-Seq procedure was performed (Figure 1A). We analyzed the transcriptomes of 700 individual cells per sample, preserving only cells with a minimum of 10 UMIs (Unique Molecular Identifier) (Supplementary Table 1). Because of the high sequence similarity between *var* genes, we applied strict mapping criteria and disallowed any multimapping during alignment.

**Figure 1.**
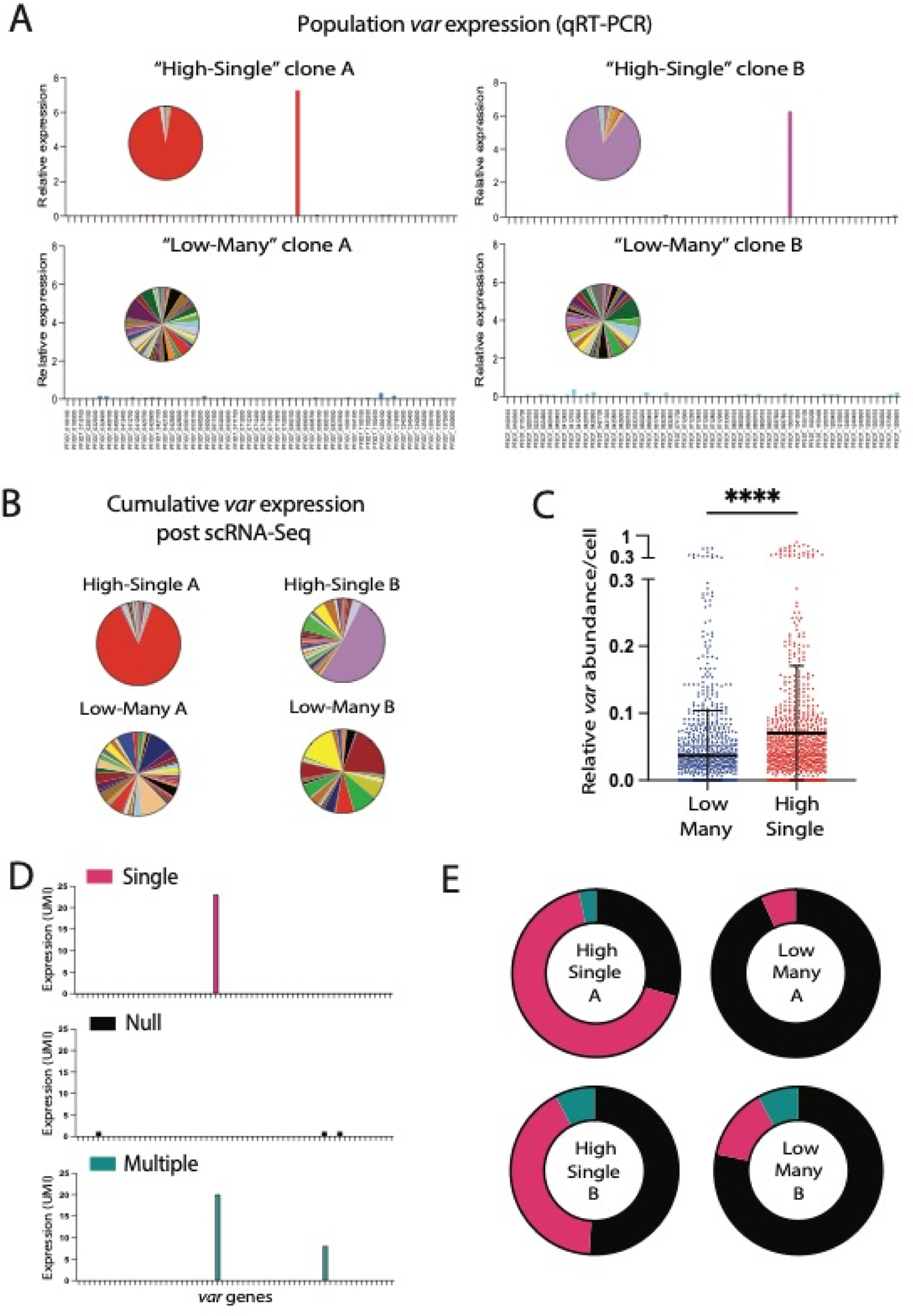
Single-cell analysis by Drop-Seq reveals different *var* expression states. (A) *var* expression profiles of the four populations analyzed in the Drop-Seq experiments, represented as histograms and pie charts. Expression of each gene is determined by quantitative RT-PCR and is represented as relative to seryl-tRNA synthetase (PF3D7_0717700). (B) Cumulative *var* expression of the four populations resulting from Drop-Seq represented as pie charts. The size of each slice of the pie is proportional to the number of UMI detected per individual *var* gene. (C) Number of total *var* UMIs relative to total UMI per individual cell in the two “low-many” populations (blue, N=1048) compared to the two “high-single” populations (red, n=873). The mean ± standard deviation is shown, and an unpaired t-test indicates a ****p < 0.0001. (D) Representative examples of individual cells in three *var* states: Single (pink), cells expressing a single *var* gene at a high-level; Null (black), cells not expressing any or a very low-level of *var* transcripts; Multiple (Blue), cells expressing two or more *var* genes at the same time. Expression is shown as number of UMI detected per individual gene. (E) Percentage of individual cells in the Single state (pink), Null state (black) or Multiple (blue) in each of the populations. Genes were considered expressed with at least 2 UMIs.

Initially, to determine the reliability of the single-cell technology for detecting *var* gene expression, we combined the individual *var* expression profiles obtained from the Drop-Seq protocol and compared the resulting cumulative expression pattern to the patterns obtained by qRT-PCR from the same populations prior to single cell isolation. In both “high-single” and “low-many” samples, the cumulative single-cell transcriptome recapitulated a similar profile to what was observed by qRT-PCR (Figure 1B), indicating that despite using very different methodologies, both techniques provide comparable assessments of *var* expression patterns. When we explored the relative number of *var* transcripts detected in individual cells we observed that cells obtained from the “high-single” populations had significantly more *var* transcripts than the parasites from the “low-many” populations (Figure 1C). More strikingly, while there was at least one detectable *var* transcript in majority of the parasites from the “high-single” populations, greater than 75% of transcriptomes from the “low-many” populations contained no detectable *var* transcripts. This suggests that majority of the individual cells in the “low-many” clones are silent, or null, from the point of view of *var* expression. In addition, in both the “high-single” and “low-many” populations, we observed that approximately 3-5% of individual parasites expressed a second *var* gene in addition to the dominant gene (Figure 1D, E, Supplementary Table 2). We refer to these parasites as “multiple”. The doublet rate (See Methods) and the possible combinations of *var* genes detected in different cells make it highly improbable for the “multiples” to be explained by RBCs infected with two parasites each expressing a different *var* gene or by doublets in the Drop-Seq procedure. For example, all “multiple” cells displayed expression of the dominant *var* gene along with a second gene, while no individual cells were detected expressing the second *var* gene alone, as would be expected if “multiples” were the result of two “single” parasites captured in the same droplet.

Overall, we could detect cells in three different *var* expression states, two of which violate the paradigm of constant and mutually exclusive expression (Figure 1D). In the “high-single” populations, the majority of individual cells displayed expression of a single *var* gene, a result that was expected and that is consistent with the standard model that each parasite expresses one and only one *var* gene at a time. In contrast, both the parasites expressing more than one *var* gene (the “multiple” cells) and those not expressing any detectable *var* genes (the “null” cells) were unexpected and are inconsistent with current models of *var* gene expression. We hypothesize that the parasites expressing more than one *var* gene might be actively undergoing a switching event, something that has not been possible to observe previously. The parasites not expressing any *var* genes are more puzzling but could represent the *var* non-expressing parasites previously observed in chronic asymptomatic infections (19). Taken together, these experiments clearly demonstrate that *var* expression is more flexible that previously thought and that, at least in cultured parasites, it is not uncommon for *var* gene expression to be extremely low or silent.

### Decreased S-adenosylmethionine synthetase activity disrupts mutually exclusive expression in individual cells

Both “multiple” cells and “null” cells were unexpected based on previous assumptions regarding mutually exclusive expression, so we chose to investigate these two states in more detail. In a previous study, we found that the availability of intracellular S-adenosylmethionine (SAM), the principal methyl donor for methylation modifications, can influence *var* expression (39). Genetic modifications to SAM synthetase (SAMS), the enzyme producing SAM from methionine, led to profound changes in *var* expression. In particular, reducing SAMS expression (40) produced populations of parasites with a total *var* expression 100-fold higher than wildtype parasites (Figure 2A). Moreover, recently cloned SAMS-KD populations displayed high expression of multiple *var* genes (Figure 2B, C), unlike wildtype parasites that typically display expression of a single dominant *var* gene (34). To gain further insights on how methylation levels influence *var* expression at the individual cell level, we analyzed two SAMS-KD clones using Drop-Seq.

**Figure 2.**
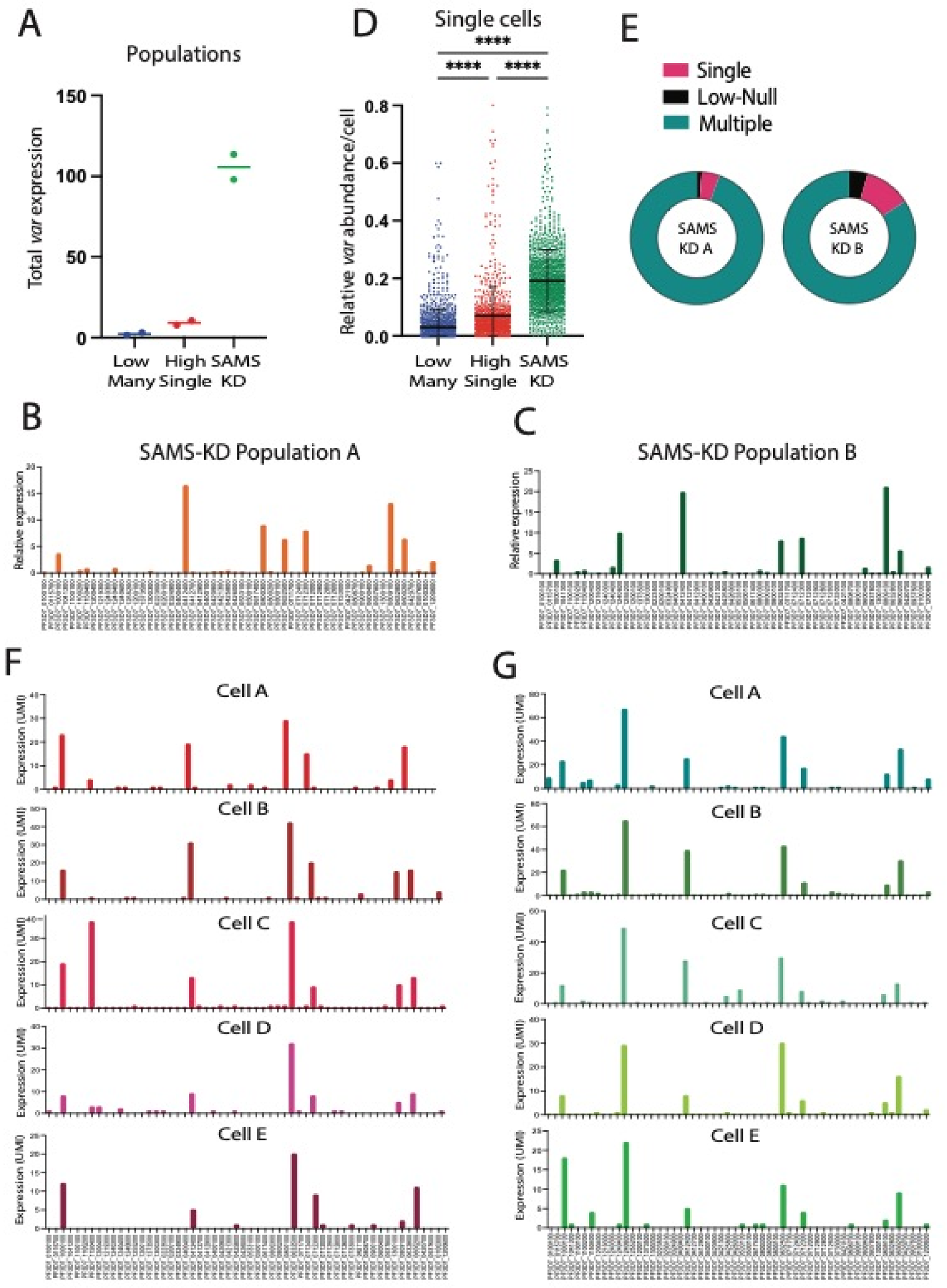
Decreased histone methylation disrupts mutually exclusive *var* gene expression in individual cells. (A) Total *var* expression levels as determined by quantitative real-time RT-PCR (qRT-PCR) for two “low-many” populations (blue), two “high-single” populations (red) and two SAMS-KD populations (green). The horizontal line shows median value. (B,C) *var* expression profiles of the SAMS-KD Population A (B) and Population B (C) determined by quantitative RT-PCR and represented as relative to seryl-tRNA synthetase (PF3D7_0717700). (D) Number of total *var* UMI relative to total UMI per individual cell in the two “low-many” populations (blue, n=1048) compared to the two “high-single” populations (red, n=873) and the two SAMS-KD populations (green, n=1199). The mean ± standard deviation is shown, and a one-way ANOVA test indicates a ****p < 0.0001. (E) Percentage of individual cells in the Single state (pink), Low-Null state (black) or Multiple (blue) in each of the SAMS-KD populations. Genes were considered expressed with at least 2 UMIs. (F, G) *var* expression profiles of 5 individual cells from Population A (F) and Population B (G) determined by Drop-Seq and represented as number of UMIs.

Similar to observations at the population level, the relative amount of *var* transcripts in individual cells is significantly higher in the SAMS-KD lines compared to wildtype parasites in both the “low-many” and “high-single” states (Figure 2D). Moreover, investigation at the single-cell level confirmed that the reduction of SAMS levels causes a disruption of mutually exclusive expression, as over 80% of individual parasites express several *var* genes at high levels (Figure E, Supplementary Table 2). Interestingly, the pattern of expression in each cell is very similar and resembles the pattern of expression in the cumulative population, with the same handful of genes being expressed in each individual cell (Figure 2F, G). There is no clear correlation in terms of chromosome position or sequence similarity between the active genes, but it appears that these genes are more prone to activation compared to others when overall methylation is reduced. These data demonstrate, for the first time, the critical role of methylation in the control of mutually exclusive expression at the single-parasite level and reveal a hierarchy of *var* gene activation, with some genes more readily activated than others.

### *var*-enrichment probes allow deeper *var* transcript detection in scRNA-Seq

As previously mentioned, another well-studied example of mutually exclusive expression is the olfactory receptor gene (*OR*) family of mammals (24). scRNA-Seq analysis of individual olfactory neurons during their maturation has shown that, before committing to expression of a single olfactory receptor gene, each neuron undergoes a developmental phase in which they express several *OR* genes at very low levels (30, 41). As the cells continue to mature, expression is narrowed to a single, highly transcribed gene. African trypanosomes were also recently shown to undergo transition from expression of many metacyclic variant surface glycoprotein (*mvsg*) genes to a single gene as they differentiate into the metacyclic form in the salivary glands of the tsetse fly vector (42). Given the parallels between these systems and *var* gene expression in *P. falciparum*, we were curious if the individual parasites in which we observed no detectable *var* transcripts might be similarly expressing a heterogenous mix of *var* transcripts that are below the threshold of detection of Drop-Seq. To improve our ability to detect even exceptionally low levels of *var* transcripts, we used xGen Hybridization Capture (Integrated DNA Technologies) and designed enrichment probes targeting all *var* genes in the 3D7 genome. As controls, we included six genes known to be expressed within the 16-19 hpi window by all cells. We additionally included PF3D7_0304600 (circumsporozoite protein, *csp*) to check for background transcription and enrichment of nonspecific targets, as *csp* is only expressed during sporozoite development in the mosquito. The design aimed to have no more than 480bp gaps between the stop of one probe and the start of the subsequent probe. This resulted in a total of 793 probes for 69 targeted genes, with an average of 11 probes per gene (See Supplementary Table 3). These probes were used to specifically pull-down and sequence only the transcripts of interest thereby greatly increasing our sensitivity for detecting *var* gene transcripts. We started with the Drop-Seq libraries from a “high-single” and a “low-many” population (Figure 1A) and prepared enrichment libraries for sequencing using Illumina technology (Figure 3A).

**Figure 3.**
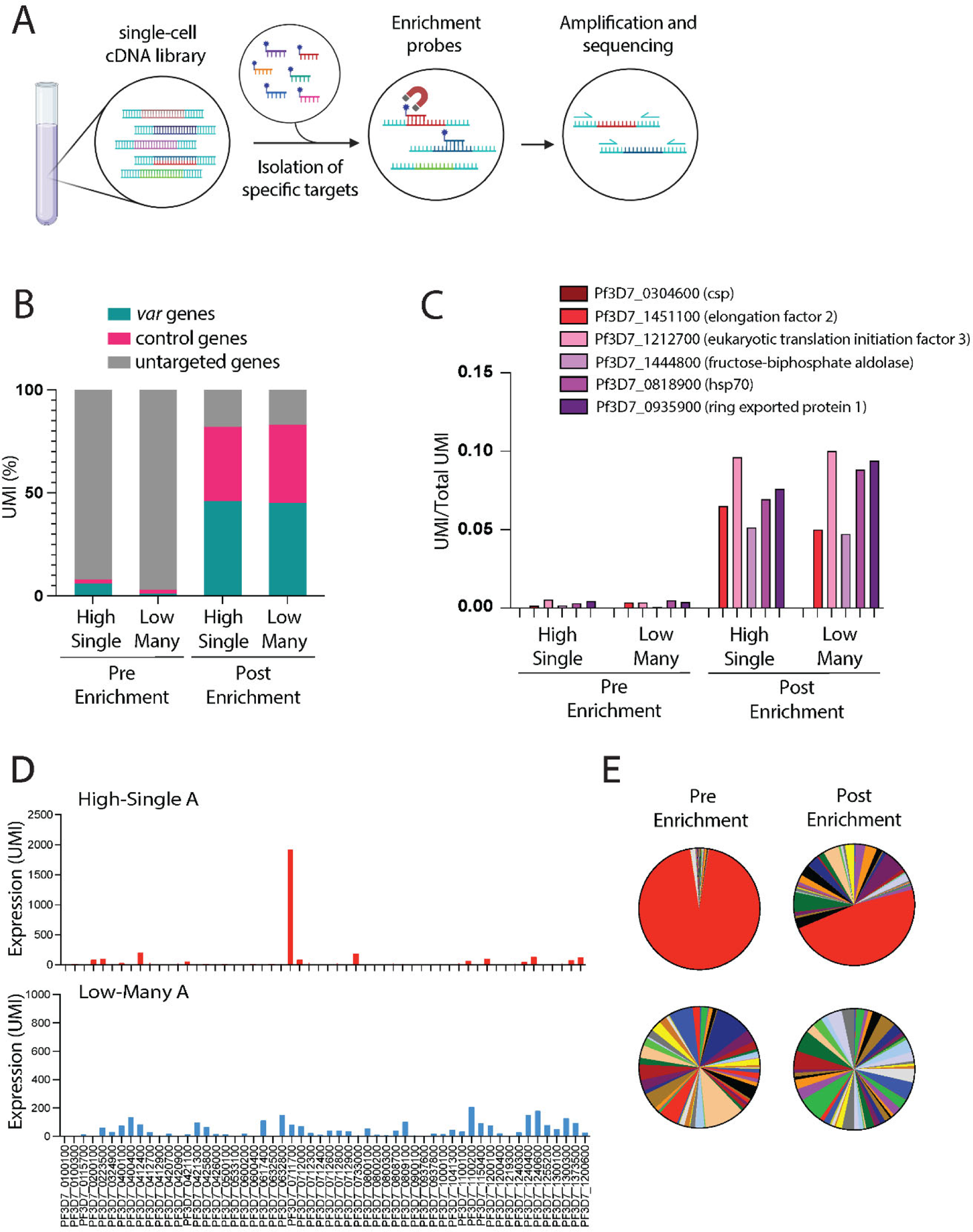
*var*-enrichment probes allow deeper *var* transcript detection in scRNA-Seq. (A) Schematic representation of the enrichment procedure. (B) UMI percentage over total UMI for *var* genes (blue), control genes (pink) and genes not targeted in the enrichment process (grey) in “high-single” and “low-many” populations before and after enrichment. (C) Number of UMI over total UMI for each control gene in “high-single” and “low-many” populations before and after enrichment. (D) *var* expression profiles after Drop-Seq and enrichment of “high-single” clone A and “low-many” clone A represented as number of UMIs. (E) Pie charts representing *var* expression profiles after Drop-Seq before and after enrichment of “high-single” clone A and “low-many” clone A.

The targeting strategy resulted in very successful enrichment for on-target transcripts. While these transcripts initially represented approximately 5% of the total transcriptome, after enrichment the on-target transcripts represent more than 80% of the total sequenced transcriptome (Figure 3B). We observed a similar enrichment of all the control genes, with the exception of the negative control *csp*, which remained undetected after enrichment (Figure 3C). To confirm that the enrichment process was not biased toward a subset of *var* genes, we combined the individual transcriptomes obtained after enrichment of the “high-single” and “low-many” populations and compared the cumulative *var* expression profiles to those obtained from the same parasite populations in the absence of enrichment. The patterns identified the same dominant *var* gene in the “high-single” population, but detected a greater amount of low expressed genes, indicative of greater sensitivity, as expected (Figure 3D, E).

When analyzed at the single cell level, *var* expression profiles from the “high-single” population were also very similar to what was observed before enrichment (compare Figure 4A to 4C and E). After enrichment, transcripts from additional genes were detectable, nonetheless a single dominant *var* gene (shown in red) was expressed at high levels in each individual cell. In contrast, for parasites in the “low-many” population, instead of most cells displaying no detectable *var* transcripts (Figure 4B), after enrichment several *var* transcripts were often detectable, indicating that these parasites are indeed expressing multiple *var* genes at very low levels (Figure 4D and F). This phenomenon resembles what was shown for olfactory receptor genes as the cells undergo the process of choosing a single *OR* gene for high level expression, suggesting the possibility that a similar mechanism of choice might be functioning in *P. falciparum*.

**Figure 4.**
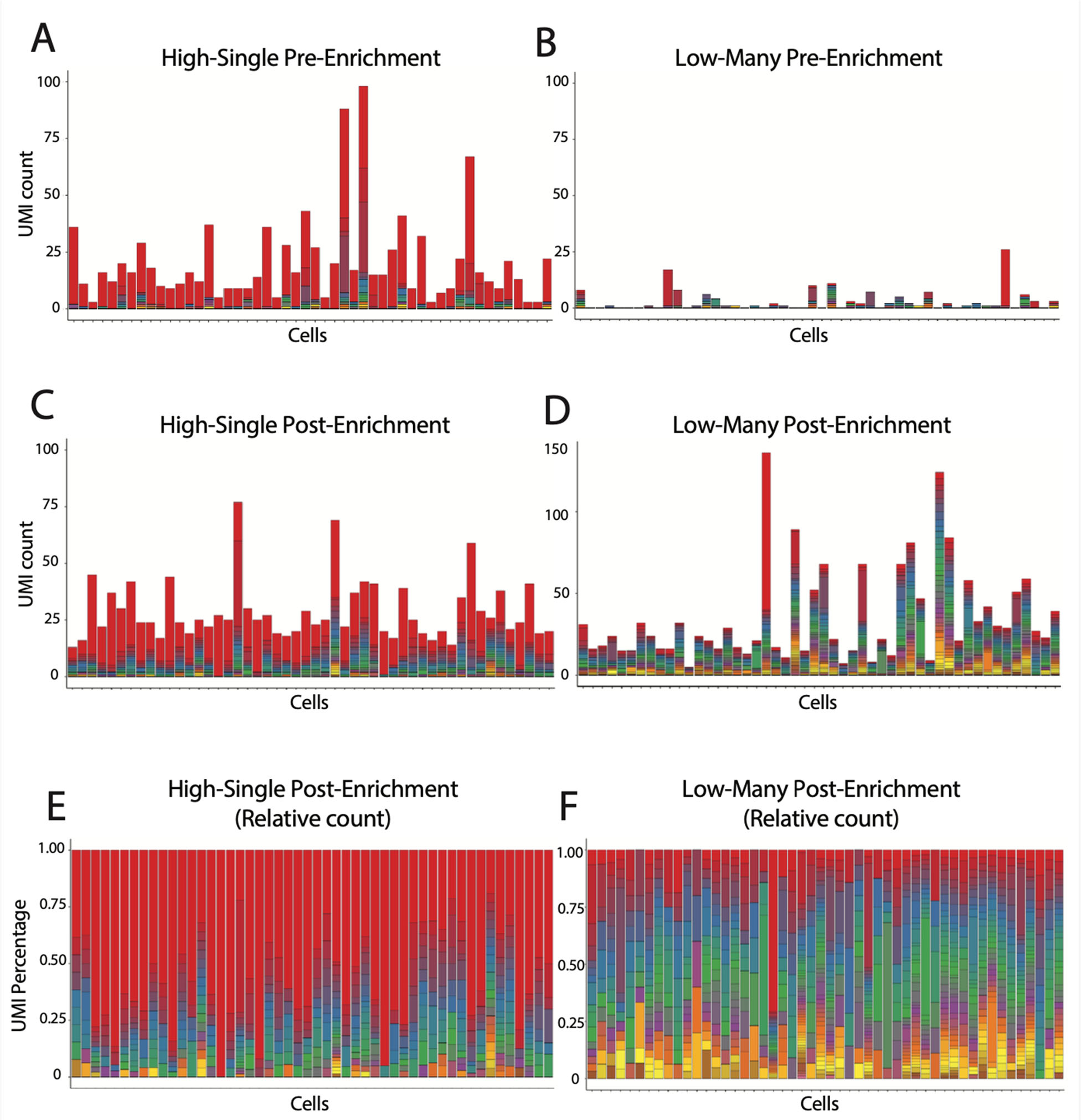
*var*-enrichment probes allow detection of multiple *var* transcripts in individual cells. *var* gene expression is displayed for the top-50 cells for “high-single” (A,C) or “low-many” (B,D) populations according to total UMI detected by Drop-Seq. *var* gene expression is shown either before (A,B) or after (C,D) enrichment. Each color in a bar represents a single *var* gene and each bar represents an individual cell. (E, F) Relative UMI counts are shown as the percentage of total *var* UMI in “high-single” (E) and “low-many” (F) from Drop-Seq after enrichment.

### A portable well-capture system for single-cell RNA-Seq confirms the different *var* expression states

To further validate the findings and address the limitations associated with Drop-Seq, we employed a portable microwell system for scRNA-Seq developed by Honeycomb Biotechnologies. As previously outlined for Drop-Seq, samples were synchronized to 16-19 hpi and enriched for infected red blood cells through SLO treatment. Infected cells were loaded and stored on HIVE devices. The procedure involves loading infected cells into a HIVE device and allowing them to gently settle into picowells containing barcoded mRNA-capture beads. The devices are then frozen and stored at −80 °C until ready for processing. Frozen devices can then be subsequently processed simultaneously for library preparation and sequencing. To enable a direct comparison to the results we obtained with Drop-Seq, we analyzed the same four populations shown in Figure 1A, applied the same strict mapping criteria and did not allow any multimapping during alignment.

The number of individual transcripts captured per cell using HIVEs was greatly improved compared to Drop-Seq. Despite loading an order of magnitude fewer cells per sample on the HIVE (1.85e^5^ cells for Drop-Seq, 1.5e^4^ cells for HIVE), and setting a quality threshold of 25 UMI per cell, we were able to recover an average of 2986 cells per sample and an average of 240 transcripts per cell (Figure 5A, B, Supplementary Table 1). The improvement in transcript capture enabled us to analyze *var* gene expression with substantially greater resolution and sensitivity. Samples underwent integration and dimensional reduction, and transcriptomes were visualized in low-dimensional space as unifold manifold approximation and projection (UMAP) plots, with the cells organized based on transcriptional similarity. Utilizing Seurat for clustering, the parasite transcriptomes clustered predominantly by differential *var* gene expression (Figure 5C, Supplementary Table 4).

**Figure 5.**
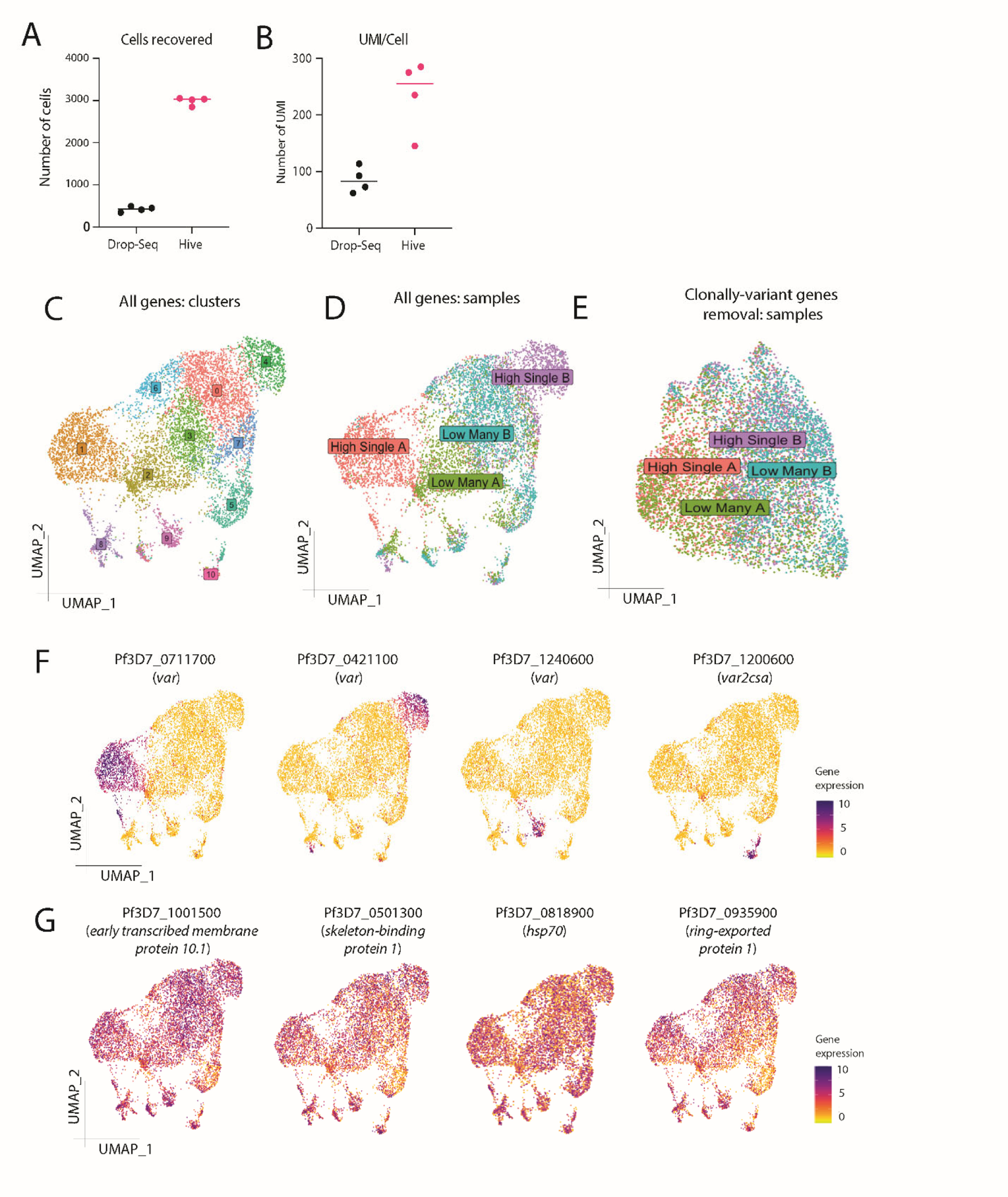
*var* genes are the main cluster-drivers in HIVE scRNA-Seq. (A) Number of cells recovered with a minimum of 25 UMI per cell in the Drop-Seq experiments (black) compared to HIVE experiments (pink). (B) Average number of UMI per cell in the Drop-Seq experiments (black) compared to HIVE experiments (pink). (C) UMAP of the HIVE single-cell transcriptomes obtained from the four parasite populations with cells colored according to their clustering (see Supplemental Table 4 for cluster information). (D) UMAP of the HIVE single-cell transcriptomes with cells colored according to the parasite population that was sampled. (E) UMAP of the HIVE single-cell transcriptomes obtained from the four parasite populations excluding clonally-variant genes from the analysis. Cells are colored according to the parasite population that was sampled. (F) UMAP graphs as in (C, D) with cells colored according to expression level of different *var* genes. (G) UMAP graphs as in (C, D) with cells colored according to expression level of different ring-expressed genes.

Cells from the two “high-single” populations distinctly clustered at opposite ends of the plot, specifically clusters 1 and 4. Notably, the key gene expression driver defining each cluster was the single dominant *var* gene expressed by the original populations, PF3D7_0711700 and PF3D7_0421100 respectively, (Figure 5D, F). In contrast, cells from the two “low-many” populations clustered in the plot’s center without significant differences in gene expression (Figure 5D). Individual parasites that had switched expression away from the dominant *var* gene were clearly identified in the smaller clusters, such as clusters 8, 9, and 10, with the alternative *var* gene differentiating each cluster (Figure 5C, F). A comprehensive list of cluster-defining genes is provided in Supplementary Table 4. The prominence of differential *var* gene expression as the primary cluster driver becomes more evident when repeating the analysis and clustering while excluding all genes known to be subject to clonally variant expression (13). In this scenario, all clustering dissipated, and individual cells from different samples are mixed with each other, indicating no discernible differences between samples except for clonally variant genes (Figure 5E). Additionally, genes known to be expressed in the 16-19 hpi window exhibited homogeneous expression across cells from different samples (Figure 5G), indicating that all samples were harvested at the same point in the replicative cycle.

These experiments confirmed the results obtained by Drop-Seq combined with enrichment. Cells from the “high-single” population expressed a dominant *var* gene at high levels (Figure 6A, C), whereas in cells from the “low-many” population, we detected low levels of transcripts from several different *var* genes at the same time (Figure 6B, D). *var* expression matrices for all cells is provided in Supplementary Table 5. This shows how HIVE scRNA-Seq represents a significant improvement in detection compared to Drop-Seq, as we could detect low levels of *var* expression without an additional enrichment step. The portability and flexibility of these devices, and the ability to easily store samples for lengthy periods of time and transport them long distances, makes them potentially particularly useful for field studies.

**Figure 6.**
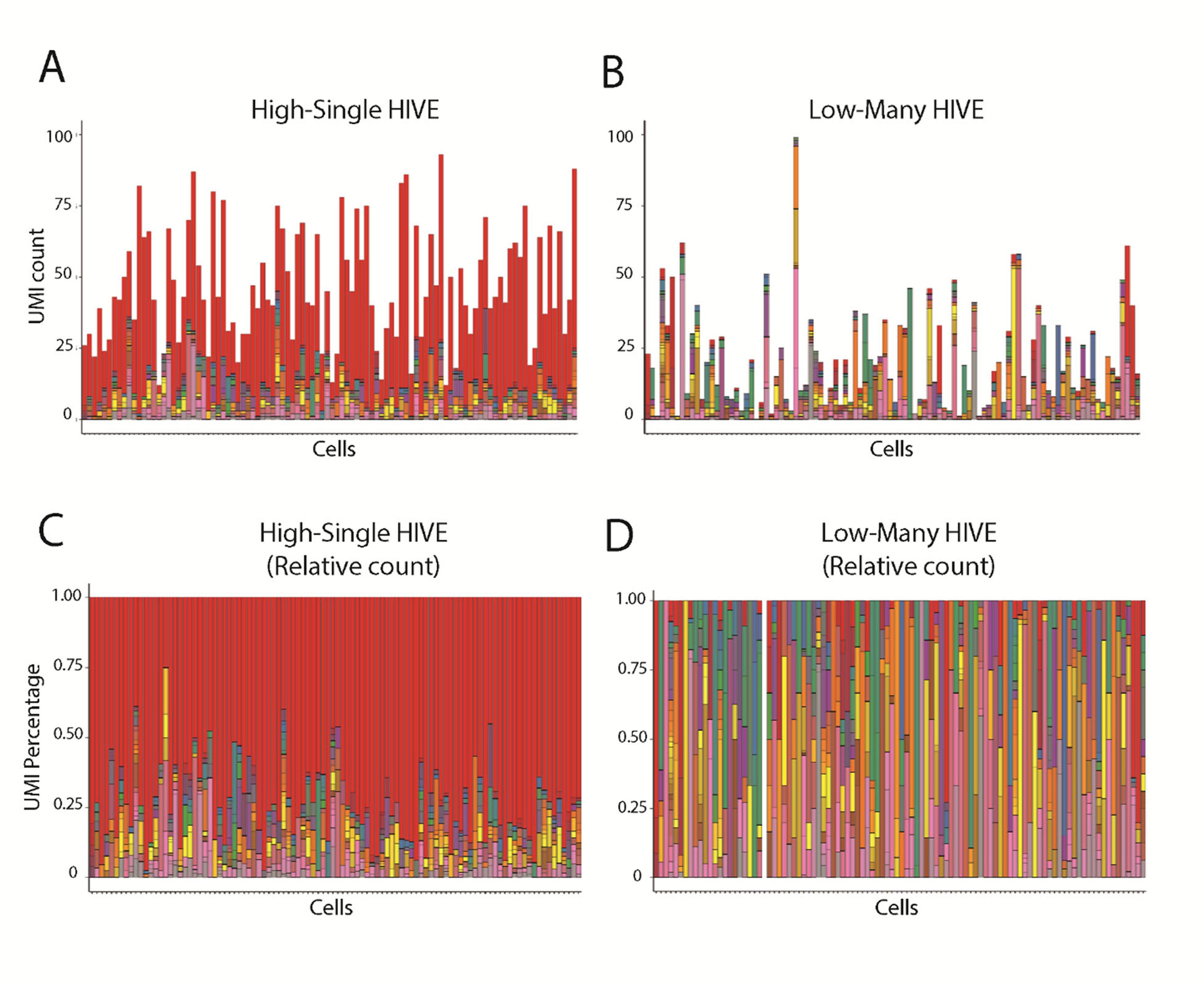
HIVE scRNA-Seq confirms multiple *var* transcripts in individual cells. *var* gene expression is displayed for the top-100 cells for “high-single” (A,C) or “low-many” (B,D) populations according to total UMI detected by HIVE scRNA-Seq. Each color in a bar represents a single *var* gene and each bar represents an individual cell. UMI counts are displayed for individual cells obtained from “high-single” (A) and “low-many” (B) populations. Relative UMI counts are shown as percentage of total *var* UMI in cells obtained from “high-single” (C) and “low-many” (D) populations.

### Greatly reduced antibody recognition of parasites in the “low-many” state

The identification of parasites expressing low levels of *var* genes raises a fundamental question: do these parasites export a detectable amount of PfEMP1 to the infected cell surface, and what implications does this have for recognition by the human immune system? If parasites in the “low-many” state indeed correspond to parasites observed in prolonged asymptomatic infections (18, 19), we anticipate their ability to better evade antibody recognition, thereby facilitating the maintenance of long-term infections.

To assess the surface expression of PfEMP1 and the corresponding immunogenicity of various parasite lines, we used pooled hyperimmune IgG obtained from 834 Malawian adults infected with *P. falciparum* (43). This IgG mixture was obtained from people who, through a lifetime of exposure to malaria infections, are largely immune to symptomatic malaria through the acquisition of antibodies capable of recognizing a broad range of Plasmodium surface antigens. Given that PfEMP1 is established as the primary target of humoral immunity (9), this reagent can be used to detect PfEMP1 surface expression (44). We used flow cytometry to measure the reactivity levels of different clonal lines using the pooled hyperimmune IgG. When initially tested against the 3D7 and NF54 strains, regardless of their *var* expression profile at the mRNA level, we observed either no or very limited reactivity (Supplementary Figure 3). Given that long-term in vitro cultivation of parasites is known to alter and reduce cytoadhesion and surface expression of PfEMP1 (45, 46), we opted to examine clonal lines that we recently generated from IT4 (Supplementary Figure 1), a parasite line known to maintain strong cytoadhesive properties and kindly provided by Dr. Joseph Smith.

We selected eight IT4 clonal populations with varying levels of cumulative *var* expression (Figure 7A) and assessed their reactivity to the hyperimmune IgG. Remarkably, a notable shift in IgG recognition was observed by parasites in the “high-single” state compared to parasites in the “low-many” state, with the latter exhibiting recognition levels similar to uninfected red blood cells (uRBC) (Figure 7B). A clear correlation of IgG recognition and the level of *var* expression is evident across all seven clones (Figure 7C), supporting the hypothesis that parasites with a low level of *var* expression remain immunologically silent. Collectively, these results support the notion that individual parasites can exist with variable levels and numbers of expressed *var* genes. Specifically, they can exist in a state characterized by very low expression of multiple *var* genes, lacking PfEMP1 on the infected RBC surface. This expression state could enable parasites to evade immune recognition and sustain long-term infections, consistent with the non-*var* expressing parasites previously observed in a chronic infection (19).

**Figure 7.**
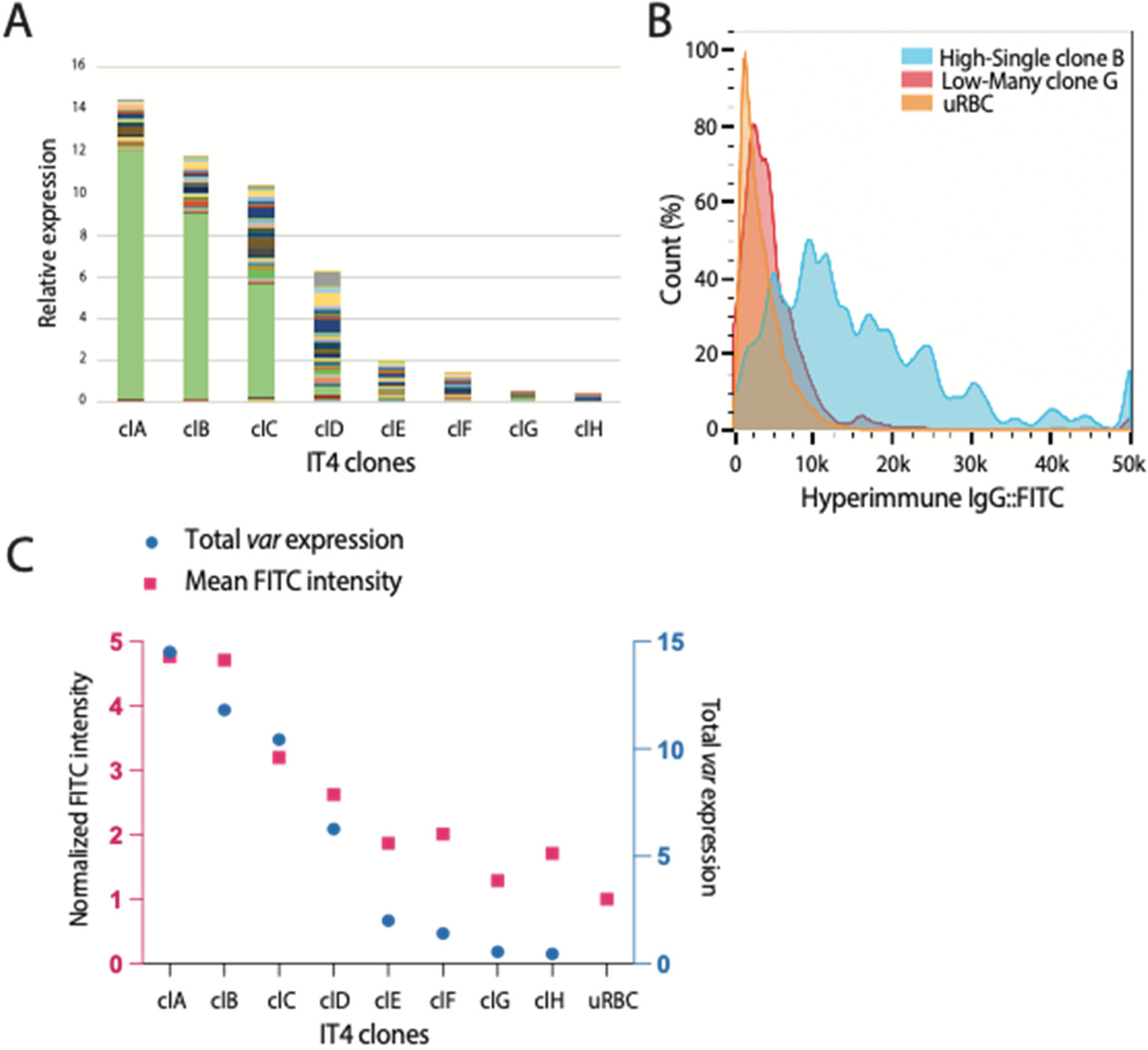
Parasites in the “low-many” state exhibit reduced immunogenicity. (A) Total *var* expression levels for IT4 clones determined by quantitative RT-PCR, with transcripts for each *var* gene shown in a different color. Values are shown as relative to seryl-tRNA synthetase (PfIT_020011400). (B) Example of flow-cytometry with hyperimmune IgG on a “high-single” IT4 clone (blue, gated infected RBC), one “low-many” (red, gated infected RBCs) and uninfected RBCs (orange). Histogram shows normalized cell count over FITC intensity. (C) Correlation between total *var* expression determined by qRT-PCR (blue) and mean FITC intensity (pink) quantified by flow-cytometry for each IT4 clone. FITC intensity of infected RBCs is normalized to FITC intensity of uRBCs.

## Discussion

Central to *P. falciparum’s* ability to sustain prolonged infections is its capacity to balance evading splenic clearance with being recognized by the immune system. Over the course of an untreated infection, parasites are thought to eventually exhaust their repertoire of *var* genes, leading to clearance of the infection (47). While this model provides an explanation for the course of a symptomatic infection, it does not fully explain the nature of asymptomatic infections that can last for years. For example, Ashley and White recently catalogued dozens of validated cases of asymptomatic infections lasting for up to 13 years (18). These infections remained untreated and unknown until revealed by splenectomy or through blood donation for transfusion. Given that PfEMP1 displayed on the RBC surface appears to readily give rise to antibodies that can clear infected cells, it is possible that once the *var* repertoire was exhausted, the infections consisted of parasites that were no longer expressing PfEMP1, similar to the IT4 parasites we examined that were not recognized by hyperimmune IgG (Figure 7). This hypothesis also provides an explanation for the rapid expansion of the parasite population after splenectomy. Consistent with this hypothesis, a study examining asymptomatic infections implicated non-PfEMP1 antigens as the primary parasite proteins on the infected RBC surface when high titers of anti-PfEMP1 antibodies are present (48), and a previous study of Kenyan children observed that parasites can reduce *var* gene expression in response to host antibodies (49). The clearest supporting evidence for this hypothesis was provided by Bachmann et al (19) who examined parasites isolated from a patient in Germany who displayed no signs of infection until after splenectomy, at which time parasitemia rose rapidly. Parasites isolated from this individual expressed no detectable *var* transcripts and did not display any cytoadhesive properties, consistent with parasites that were not expressing PfEMP1. Taken together, these studies suggest a model in which parasites face two intense and opposing selective pressures: 1) clearance by the spleen and 2) recognition by anti-PfEMP1 antibodies. In the absence of high titers of anti-PfEMP1 antibodies, PfEMP1-expressing parasites rapidly expand, leading to high parasitemias, severe illness and the waves of parasitemia typically observed in symptomatic *P. falciparum* infections. However, in individuals who have significant anti-PfEMP1 immunity, PfEMP1 expression is selected against, enabling only “PfEMP1-null” parasites to survive. Due to splenic clearance, these infections remain at a low parasitemia and cause little or no symptoms, resulting in chronic infections that could last indefinitely. However, it is worth noting that such infections could potentially contribute to transmission of the disease. For example, one study recently attributed the bulk of transmission to asymptomatic infections (50), and since asymptomatic individuals seldom seek treatment, this potential parasite reservoir could contribute substantially to disease transmission and complicate efforts to eliminate or eradicate malaria.

This hypothesis predicts that “PfEMP1-null” parasites will dominate long-term infections where antibodies have eliminated PfEMP1-expressing parasites. This would provide an explanation for the asymptomatic nature of chronic infections and the observation of elevated parasite loads within the spleens of asymptomatically infected individuals (51). Further, it also offers an explanation for instances of individuals who, after residing outside endemic regions for several years, experienced malaria relapses following splenectomy (18, 19) or during pregnancy (52–55). Unlike other *Plasmodium* species that can form hypnozoites (56), *P. falciparum* does not naturally have a dormant state, and therefore requires an alternative strategy to maintain prolonged infections and bridge periods with low or zero transmission. It has been observed that parasites present at the end of a dry season or in chronic asymptomatic infections display decreased cytoadherence, with increased splenic clearance and lower parasitemias (57, 58), properties consistent with the “PfEMP1-null” state. This could provide an explanation for how *P. falciparum* can persist within a geographical region for prolonged periods in the absence of transmission without entering a dormant state. An intriguing question remains regarding whether these parasites persist solely in circulation at extremely low levels or potentially sequestered in a specific location, as observed for gametocytes in the bone marrow (59).

Our observation that individual parasites from populations in the “low-many” state are actually expressing many *var* genes at a very low level is reminiscent of OR gene expression in maturing olfactory neurons during the process of single gene choice (30, 41). For OR gene expression it has been speculated that this represents a developmental state in which promoters are competing for activation, after which a single gene becomes dominantly expressed. The concept of promoter competition could provide an explanation for the different *var* gene expression states that we detected. For example, in addition to parasites expressing very low levels of *var* transcripts, we also observed parasites expressing high levels of *var* transcripts from multiple genes, a phenotype that was greatly enhanced through reduced expression of PfSAMS (Figure 2). We hypothesize that when SAM availability is reduced, histone methylation is similarly lowered, enabling more than one *var* gene to compete for activation. This has implications for mechanisms that could affect *var* expression switching, for example conditions that reduce SAM availability could loosen mutually exclusive expression and thus enhance promoter competition, leading to expression switching. Interestingly, our single cell analysis of *var* gene expression in the PfSAMS knockdown lines detected activation of a specific subset of *var* genes, consistent with the hypothesis that certain *var* genes are more prone to activation than others. This is similar to our previous observations that *var* activation is biased toward certain subsets of *var* genes and that this bias shifts overtime, potentially shaping the trajectory of *var* gene expression over the course of an infection (34). What characteristics determine the likelihood of any particular gene becoming activated remain unknown, although methylation appears to be key to limiting the number of active genes.

## Acknowledgments

The authors would like to thank Drs. Kami Kim and Alister Craig for providing access to hyperimmune IgG and Dr. Joseph Smith for providing the IT4 parasite line. This work was supported by the National Institutes of Health (AI 52390 and AI99327 to K.W.D). K.W.D. is a Stavros S. Niarchos Scholar and a recipient of a William Randolf Hearst Endowed Faculty Fellowship. F.F. received support from the Swiss NSF (Early Postdoc.Mobility grant P2BEP3_191777). J.E.V. received support from F31 Predoctoral Fellowship F31AI164897 from the NIH. The Department of Microbiology and Immunology at Weill Medical College of Cornell University acknowledges the support of the William Randolph Hearst Foundation. The funders had no role in the study design, data collection and analysis, decision to publish, or preparation of the manuscript.

## Author contributions

F.F., J.E.V., E.H. and S.M. designed and performed the experiments, collected, and analyzed the data. B.F.C.K. aided in designing custom scripts for data analysis. F.F., J.E.V., E.H., C.N., B.F.C.K. and K.W.D. wrote the paper.

## Methods

### Parasites culture

Both 3D7 and IT4 parasite lines were maintained following standard procedures at 5% hematocrit in RPMI 1640 medium supplemented with 0.5% Albumax II (Invitrogen) in an atmosphere containing 5% oxygen, 5% carbon dioxide, and 90% nitrogen at 37°C. Clonal parasites lines were obtained by limiting dilution in 96-well plates, with an average of 0.5 parasites per well (60).

### RNA extraction, cDNA synthesis and RT-qPCR

For RT-qPCR, RNA was extracted from ring-stage parasites 48 hours after synchronization with 5% Sorbitol (61). RNA was extracted with TRiZol (Invitrogen) and purified on PureLink (Invitrogen) columns following the manufacturer’s protocols. To eliminate genomic DNA, RNA was treated with DNase I (Invitrogen). cDNA was synthesized from 1μg of RNA with Super Script II Reverse Transcriptase (Invitrogen), according to manufacturer instructions. Established sets of *var* primers for either 3D7 (62) or IT4 (63) were employed to determine *var* transcription through RT-qPCR. All reactions were performed in 10μl volumes in 384-well plates using iTaq Universal SYBR Green Supermix (Bio-Rad) in a QuantStudio 6 Flex (ThermoFisher). ΔCT for each primer pair was determined by subtracting the individual CT value from the CT value of seryl-tRNA synthetase (PF3D7_0717700) and converting to relative copy numbers with the formula 2^ΔCT^. Relative copy numbers are plotted using GraphPad Prism 10 or Microsoft Excel as bar graphs and pie charts.

### Preparation of parasites for single-cell RNA-Seq

Ring-stage cultures were initially synchronized using 5% Sorbitol. Approximately 24 hours later, late-stage parasites were isolated using percoll/sorbitol gradient centrifugation (64, 65) and allowed to reinvade for 3 hours while shaking continuously to minimize multiple infections of individual RBCs. 0-3 hours ring-stage parasites were isolated through a second percoll/sorbitol centrifugation and allowed to progress to 16-19 hpi when infected RBC were enriched by treatment with Streptolysin-O (SLO), as previously described (66). Each of this process step was verified by microscopy and cultures were only used for scRNA-seq if multiply-infected RBCs (RBCs) comprised less than 5% of all infected cells. Following enrichment for infected cells, parasites were suspended in PBS (supplemented with 0.01% BSA if preparing for Drop-Seq). Parasites were adjusted to the desired target densities of 1.85×10^5^ cells per mL for Drop-Seq and 1.5×10^4^ cells per mL for HIVE.

### Drop-Seq

Drop-Seq single-cell transcriptomes were generated as previously described (38). Briefly, droplets were generated using customed microfluidic devices (see CAD file from http://mccarrolllab.com/dropseq/, manufactured by FlowJEM). Droplets are composed by individual cells dissolved in lysis buffer and uniquely barcoded beads (ChemGenes, as designed by Macosko et al, 2015 (67)). Flow rates, beads concentration and doublet rates were based on previous optimization by Poran and Nötzel, 2017 (38). Droplets were disrupted and reverse transcription was performed with template switching to allow for cDNA amplification by PCR. 3000 beads were used for 30 cycles of PCR amplification with TSO primer (Template Switch Oligo:AAGCAGTGGTATCAACGCAGAGT). cDNA libraries were purified using Agencourt AMpure XP (Beckman Coulter). The quality of the libraries was assessed by High Sensitivity D5000 ScreenTape (Agilent Technologies). Samples were prepared for sequencing using the Nextera XT kit (Illumina), followed by two AMpure purifications at 0.6X ratio and 1X ratio respectively. Library pools concentrations were measured using Qubit dsDNA HS Assay Kit (Thermo Fisher Scientific) and High Sensitivity D5000 ScreenTape (Agilent Technologies), and subsequently sequenced on Illumina NextSeq500 with custom primers according to the original protocol (38).

### Drop-Seq transcriptome analysis

Raw reads were processed and aligned using STAR aligner (version 2.7.10a) using the standard Drop-seq pipeline, and according to the ‘Drop-seq Alignment Cookbook’, both found at http://mccarrolllab.com/dropseq/. Reads were aligned to *Plasmodium falciparum* 3D7 transcriptome (PlasmoDB v.32)(68). Considering the sequence similarity between *var* genes, for each read only a single mapping position was retained and all ambiguously mapping reads were discarded. Known non-poly-adenylated transcripts were discarded before analysis. Expression matrices were generated using cell barcodes and unique molecular identifiers (UMI). Cells were filtered and discarded if they contained genes detected in less than 3 cells or contained less than 10 UMI. Data Normalization and differential expression analysis were performed using the Seurat R package (version 4.1.0) (69).

### Enrichment

Probes design was performed in collaboration with Integrated DNA Technologies (IDT). Probes were subjected to off-target QC, where hits were counted if the match represented 90% identity over 50% of the probe length, using release 57 of *P. falciparum* 3D7 genome (PlasmoDB). Probes tiling was set to 0.2X, aiming to have no more than 480bp gaps between the stop of the previous probe and the start of the subsequent probe, resulting in 793 probes (See Supplementary Table 3). Enrichment was done according to the manufacturer protocol (xGen hybridization capture of DNA libraries, IDT). Briefly, 1 μg of Drop-Seq full-length cDNAs before Illumina Tagmentation was used as template. TSO and polyT oligos were employed for blocking and mixture were dried-down for 40 minutes at 45C using a SpeedVac system. Hybridization incubation was performed at 65C for 16 hours. Washes and streptavidin capture was done according to the manufacturer. The TSO primer was used for 12 cycles of post-capture PCR, followed by purification with AMpure beads. Samples were prepared for sequencing using the Nextera XT kit, as described above for Drop-Seq. Library pools concentrations were measured using Qubit dsDNA HS Assay Kit and the quality of the libraries was assessed by High Sensitivity D5000 ScreenTape (Agilent Technologies), followed by sequencing using Illumina NextSeq500.

### Hive

Parasites were prepared for HIVE scRNA-Seq as described above and diluted to 1.5×10^4^/ml in PBS prior loading on the device. scRNA-Seq was performed using HIVE scRNA-Seq v1 (Honeycomb Biotechnologies). Sample capture was performed according to the manufacturer’s protocol. Briefly, the HIVE device was thawed at RT for one-hour prior loading, and 15,000 cells were loaded per HIVE. Parasites were deposited in the wells via centrifugation and cells were stored in Cell Preservation Solution. Individual HIVEs were stored at −20C and batch-processed together. HIVE processing was performed according to the manufacturer’s instructions (Honeycomb Biotechnologies). All HIVE devices were processed to cDNA in a single batch according to manufacturer’s instructions (v1 revision A). Final library concentrations were measured using Qubit dsDNA HS Assay Kit (Thermo Fisher Scientific) and profiled by High Sensitivity D5000 ScreenTape (Agilent Technologies). Sequencing was performed on Illumina NovaSeq (Illumina) with custom primers (Honeycomb Biotechnologies).

### HIVE transcriptome analysis

The HIVE BeeNet pipeline (v1.1, Honeycomb Biotechnologies) was used to process the raw data into count matrices. No multimapping was allowed, using the STAR argument - -star-args=’--outFilterMultimapNmax 1’. The Seurat R package (69) was used for all the downstream analysis on the count matrices. A Seurat object was created for each of the samples and combined into a merged Seurat. Transcriptomes with less than 100 UMIs were discarded from downstream analysis. Seurat objects were normalized and variance stabilized using SCTransformation, and the resulting data were subjected to principal component analysis. Non-linear dimensionality reduction was performed using uniform manifold approximation and projection (UMAP) on the first 10 dimensions. For the identification of cluster-specific gene markers and differential gene expression between samples, the FindClusters (obj, resolution = 0.5) and FindAllMarkers (obj, logfc.threshold = 0.25, test.use = “wilcox”, min.pct = 0.25) functions in Seurat were used. For the analysis of clusters in Figure 5E and Supplementary Table 4, we removed all clonally-variant genes (13).

### Flow-cytometry

Ring-stage cultures were synchronized using 5% Sorbitol. Approximately 24 hours later, 400 μl of 2-3% late-stage parasites were twice washed with and resuspended in 400 µL of incomplete culture media and split into 4 tubes to include single-stained controls, treated or untreated with 500 μg of pooled human serum (43), and incubated for 60 minutes at room temperature. Cells were then washed three times, and treated with 16 μM Hoechst 33342 and/or Anti-Human IgG (Fc specific)−FITC antibody (1:100 dilution, Sigma-Aldrich) in iCM, and incubated for 30 minutes at 4C. After incubation, cells were washed three times with PBS and analyzed on Aurora flow cytometer with SpectroFlo (Cytek Biosciences). Flow-cytometry data were analyzed using FlowJo v10 software and GraphPad Prism 10.

### Materials Availability

All unique/stable reagents generated in this study are available from the Lead Contact without restriction.

### Data and Code Availability

All sequencing data produced for this study is deposited in the NCBI Sequence Read Archive available at https://www.ncbi.nlm.nih.gov/sra under the study accession code PRJNA1075333. All code utilized for analysis and figure production as well as count matrices are available on GitHub at https://github.com/DeitschLab/SingleCell.

---

**Figure S1.**
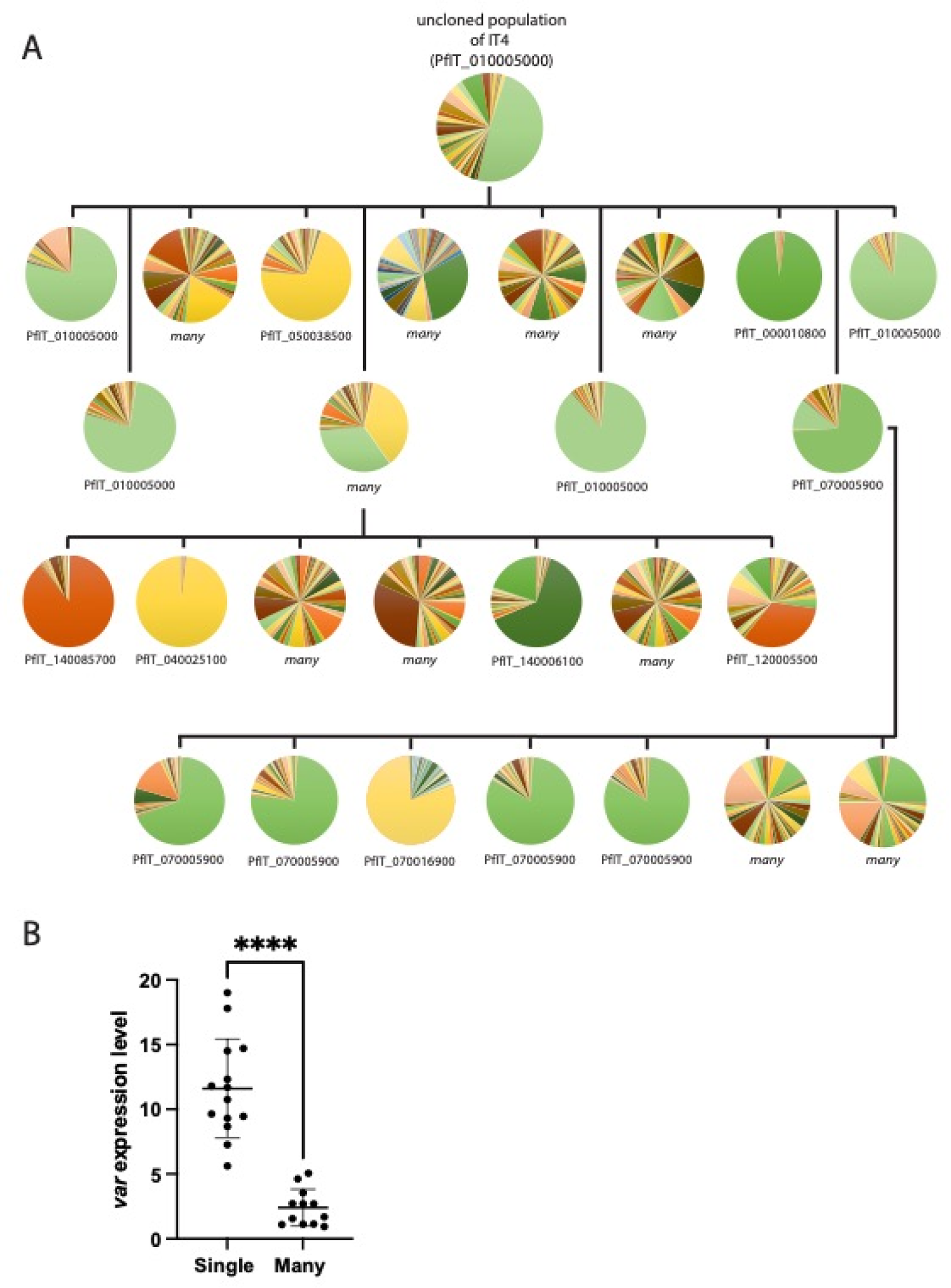
Detection of “high-single” and “low-many” *var* expression states in IT4. (A) Clone tree of wildtype IT4 parasites. Each pie chart represents the *var* profile of an individual subcloned population determined by qRT-PCR, each slice of the pie represents the expression level of a single *var* gene. The annotation number of the main *var* gene expressed is shown below for populations expressing a dominant *var* gene. Vertical and horizontal lines delineate sequential rounds of subcloning by limiting dilution. (B) Total *var* expression levels as determined by qRT-PCR for all the subclones in (A). The mean ± SD interval is shown, and an unpaired t-test indicates a ****p < 0.0001.

**Figure S2.**
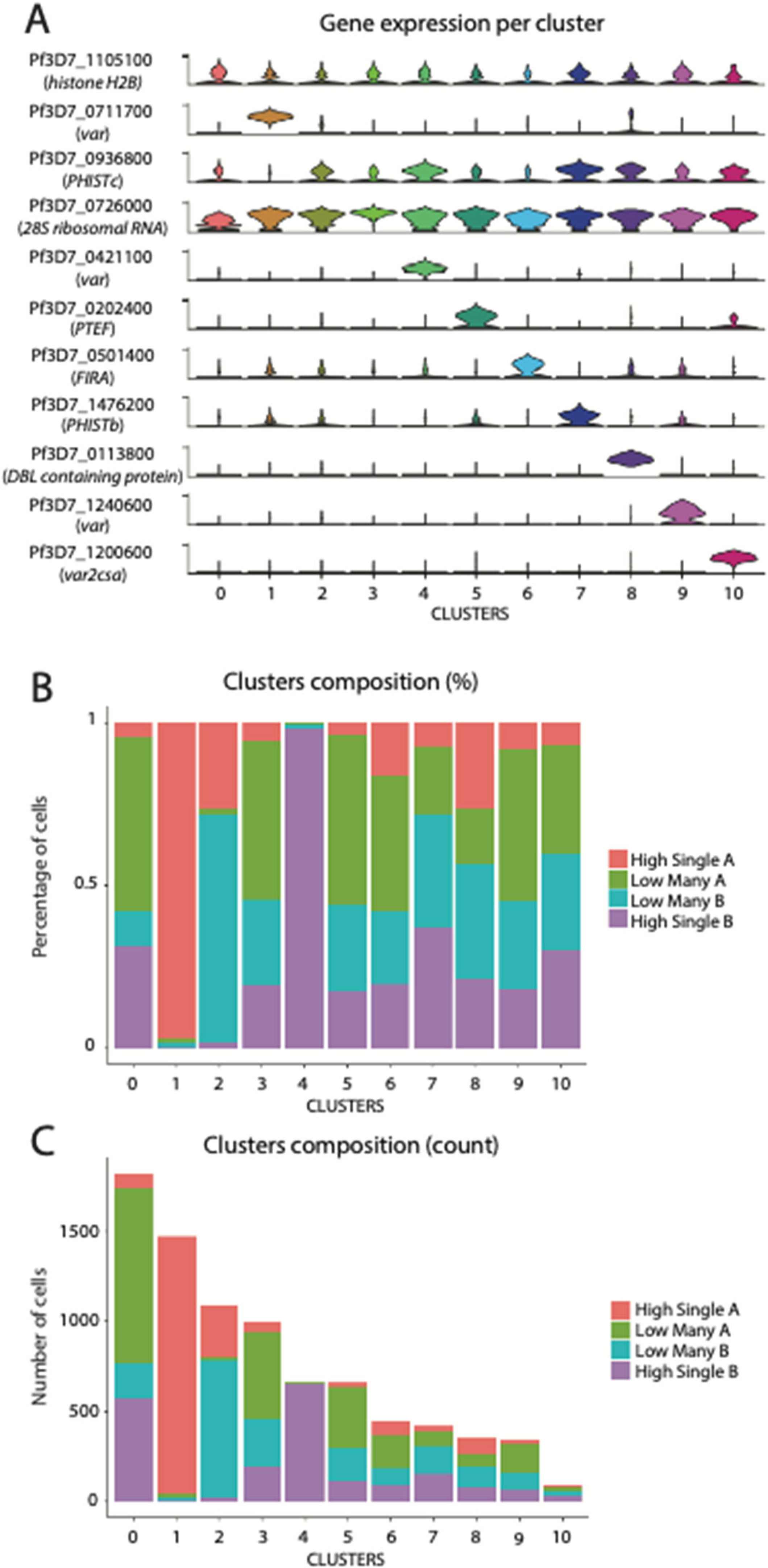
HIVE clusters composition. (A) Violin plots of gene expression per cell within each cluster in Figure 5C. Genes depicted are the most highly expressed genes in each of the clusters. (B) Percentage of cells in each cluster from Figure 5C belonging to a certain original sample. (C) Number of cells in each cluster from Figure 5C belonging to a certain original sample.

**Figure S3.**
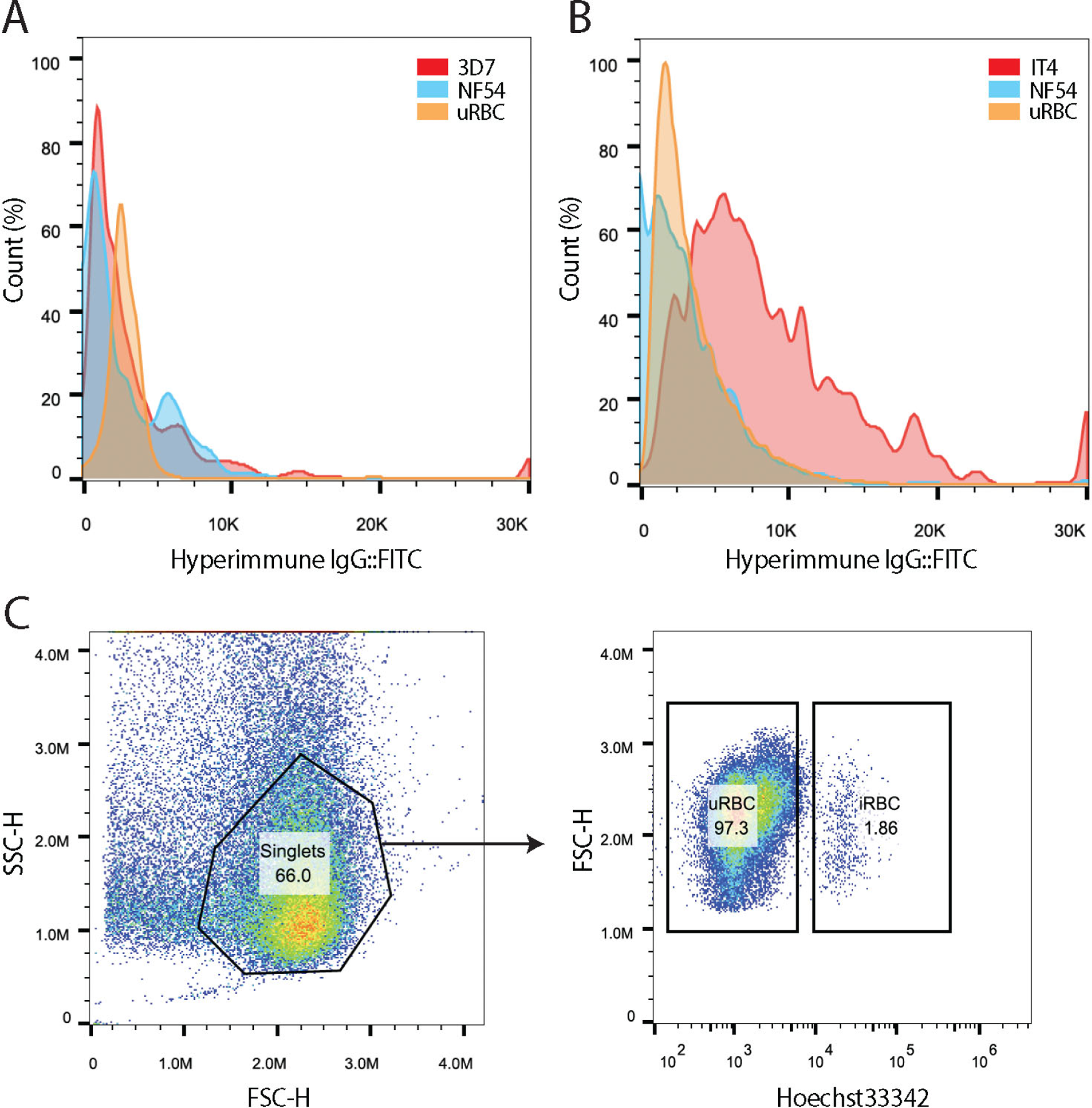
3D7 and NF54 lines exhibit low immunogenicity. (A) Flow-cytometry with hyperimmune IgG on NF54 (blue, gated infected RBC), 3D7 (red, gated infected RBCs) and uninfected RBCs (orange). (B) Flow-cytometry with hyperimmune IgG on NF54 (blue, gated infected RBC), IT4 (red, gated infected RBCs) and uninfected RBCs (orange). Histograms show normalized cell count over FITC intensity. (C) Example of flow-cytometry gating strategy applied to all experiments shown in Figure 7 and Supplementary Figure 3A and B. FSC vs SSC is initially used to identify singlets. The DNA content measured by staining with Hoechst 33342 vs FSC is used to distinguish uninfected red blood cells (uRBC) from infected red blood cells (iRBC). These gating parameters are then used directly to detect antibody recognition as displayed in the associated histograms.

